# Penile cavernous sinusoids are Prox1-positive hybrid vessels

**DOI:** 10.1101/2023.09.01.555933

**Authors:** Sarah Schnabellehner, Marle Kraft, Hans Schoofs, Henrik Ortsäter, Taija Mäkinen

## Abstract

Endothelial cells (ECs) of blood and lymphatic vessels have distinct identity markers that define their specialized functions. Recently, specialized hybrid vasculatures with both blood and lymphatic vessel-specific features have been discovered in multiple tissues. Here, we identify the penile cavernous sinusoidal (pc-S) blood vasculature as a new hybrid vascular bed expressing key lymphatic EC identity genes *Prox1, Vegfr3* and *Lyve1*. Using single cell transcriptome data of human corpus cavernosum tissue, we found heterogeneity within pc-S endothelia and observed distinct phenotypic alterations related to inflammation response in hybrid ECs in erectile dysfunction. Molecular, ultrastructural and functional studies further establish shared hybrid identity of pc-Ss in mouse, and reveal their morphological adaptations and ability to perform lymphatic-like function in draining high molecular weight tracers. Interestingly, we found that inhibition of the key lymphangiogenic growth factor VEGF-C did not block the development of pc-Ss in mice, distinguishing them from other lymphatic and hybrid vessels analyzed so far. Our findings provide a detailed molecular characterization of hybrid pc-Ss and pave the way for the identification of molecular targets for therapies in conditions of dysregulated penile vasculature, including erectile dysfunction.

## Introduction

Blood and lymphatic vessels fulfil important homeostatic functions and play a role in numerous diseases. The endothelial cells (ECs) of blood and lymphatic vessels develop during early embryogenesis and differentiate into distinct cell lineages, characterized by the expression of specific molecular markers. However, recent research shows remarkable EC plasticity that drives cell identity transitions during injury, disease and even normal physiological changes in the vasculature. In addition, organ-specific functional specialization is attributed to certain vascular beds adopting ‘mixed’ blood-lymphatic vessel identities. For example, blood ECs (BECs) of liver sinusoids and high endothelial venules of secondary lymphoid organs are characterized by a partial lymphatic EC (LEC) identity due to the expression of the key lymphatic markers LYVE1 and VEGFR3 [1]. In addition, a ‘hybrid’ blood-lymphatic endothelial phenotype was discovered in ECs of the Schlemm’s canal (SC) of the eye [2–4], the ascending vasa recta of the renal medulla [5], and the remodeled spiral arteries of the placental decidua [6]. Uniquely, these hybrid vasculatures are directly connected to blood vessels and in most cases blood perfused, yet express PROX1, which is considered the master transcriptional regulator of LEC fate [7]. The molecular properties of blood vessels with hybrid identity reflect their functional adaptation to perform ‘lymphatic-like’ functions in response to tissue demands, such as drainage of aqueous humour by the SC in the eye [1].

A remarkable example of a vascular bed that is exposed to varied environmental demands, including a range of pressure, stretch and shear forces, is the penile cavernous sinusoidal vasculature. Penile cavernous sinusoids (pc-Ss) are important for erectile functionality as they entrap blood while engorging, thereby facilitating the organ’s rigidity. The penile cavernous tissues consist of pc-Ss that are embedded within sinusoidal smooth muscle (sSM), supporting connective tissue and blood supplying and/or draining vasculature, all together enveloped by the tunica albuginea [8]. Erectile functionality depends on a complex interplay between cavernous nerves and their closely associated sSM, regulating the blood flow from the helicine arteries into the pc-Ss. During the flaccid state, the pc-Ss are tonically constricted by sSMs and only slowly perfused by blood, while when erected, the sSM and pc-Ss relax due to released neurotransmitters allowing for increased blood supply by the helicine artery and sinusoidal engorgement [9,10]. Erectile dysfunction (ED), a common condition that decreases quality of life in the male population, is presented by a wide range of psychological, physiological, as well as pathological causes. ED is a common comorbidity in other diseases, including type 1 diabetes [11]. While structural abnormalities in pc-Ss have been shown to contribute to ED [12,13], little is known about their molecular causes.

Contrary to the human penis, the mouse penis is anatomically segmented into the distal glans and the proximal body connecting at a ninety degree-angle bend of the urethra [14]. During the non-erected resting state, the glans is laying within the external preputial space which is created by the external preputial lamina terminating onto the surface of the glans. Thereby, the penis can be projected outward during the erected state. Several different, functionally specialized cavernous tissues have been identified in the mouse penis, of which those found in the penile body are considered to be analogous to the human corpus spongiosum and corpus cavernosum [15]. Additionally, cavernous rigidity is supported in mice by the baculum bone (os penis) in the glans penis [16,17].

Here we report that the blood perfused pc-Ss display molecular features of hybrid vessels and display distinct phenotypic alterations in ED. Like other hybrid vessels, they express the LEC master regulator Prox1 but, uniquely, develop independent of the key lymphangiogenic growth factor VEGF-C. Molecular characterization of pc-Ss can provide insight into their specialized functionality and identify potential therapeutic targets of the male reproductive system.

## Materials and methods

### Mice

*R26-mTmG* [18] and *R26-tdTom* [19] reporter lines were obtained from the Jackson Laboratory. *Prox1-GFP* [20], *Cldn5-GFP* [21], *Cdh5-CreER^T2^* [22] and *Vegfr3-CreER^T2^* [23] lines were described previously. All mice were maintained on a C57BL/6J genetic background. For AAV experiments, neonatal pups received a single intraperitoneal injection (10 µl) with 5×10^10^ viral particles of a recombinant AAV9 encoding the ligand binding domains 1–4 of VEGFR3 fused to an mFc domain (mVEGFR3_1-4_-mFc), or the ligand non-binding domains 4–7 of VEGFR3 fused to an mFc domain (mVEGFR3_4-7_-mFc) [24]. Experimental procedures on mice were approved by the Uppsala Animal Experiment Ethics Board (permit numbers 130/15 and 5.8.18-06383/2020) and performed in compliance with Swedish legislation.

### Tracer experiments

To visualize blood perfusion, intravenous tail vein injection with biotinylated Lycopersicon esculentum (tomato) lectin (Vector Laboratories; B-1175-1) or Lycopersicon esculentum (tomato) lectin, DyLight® 649 (Vector Laboratories; DL-1178-1) was followed by terminal anesthesia with 100 mg/kg Ketador (Richter Pharma AG) and 10 mg/kg Rompun (Bayer; 02-25-45), and cardiac perfusion fixation with 1% paraformaldehyde. Tissues were post-fixed for 1.5-2 h in 3% paraformaldehyde at room temperature and further processed for immunofluorescence analysis.

To assess fluid and macromolecular uptake by penile cavernous sinusoids, 0.5 µl of lysine-fixable tetramethylrhodamine conjugated dextran, 2 000 000 MW (Invitrogen™, D7139; 10 mg/ml in sterile filtered PBS), was injected subcutaneously in the penile glans using a Hamilton syringe (34 G). Animals were killed at the indicated timepoint by cervical dislocation. Penis and lymph nodes were dissected and fixed for 1.5-2 h in 3% paraformaldehyde (penis) or overnight in 1% paraformaldehyde (PFA, Sigma, P6148) at 4°C (lymph nodes) and further processed for immunofluorescence analysis.

### Immunoblotting

To validate AAV transduction, whole blood was collected from the thoracic cavity prior to cardiac perfusion, followed by coagulation (30 min, room temperature) and serum (supernatant) collection. Serum was purified by centrifugation (two times for 10 min, 2000xg, 4°C) and consecutive collection on ice. Bromophenol blue and 5% β-mercaptoethanol were added to the obtained serum, boiled for 95°C for 5 min and used to detect the soluble VEGFR3_1–4_-Ig or VEGFR3_4–7_-Ig serum proteins by western blotting. 50 and 100 ng recombinant mouse VEGFR3 chimera protein (R&D Systems, 743-R3-100) was used as quantitative standards. Proteins were separated by sodium dodecyl sulfate-polyacrylamide gel electrophoresis (SDS–PAGE) and blotted onto polyvinylidene difluoride membranes. Mouse VEGFR3 domains 1-4 and 4-7 were detected by probing with the polyclonal goat anti-mouse VEGFR3 antibody (R&D Systems, AF743, 1:1000) against the extracellular domain of VEGFR3. Signal detection was done using anti-goat secondary antibodies conjugated to horseradish peroxidase (0.375 µg/ml, Jackson Immuno-Research) in combination with enhanced chemiluminescence solution (Thermo-Fisher Scientific, WP20005). Visualization was performed on a ChemiDoc MP imaging system (Biorad).

### Immunofluorescence

Fixed penile tissue was decalcified in 0.5 M EDTA (pH 8) at room temperature for five days, with solution exchanged daily. The tissue was embedded in gelatin-albumin matrix and 76-90 µm coronal sections were prepared using a vibratome. The sections were permeabilized in 0.3% Triton X-100 in PBS (PBS-Tx), followed by blocking in 3% bovine serum albumin (BSA; Sigma, A3294-100G) in PBS-Tx for 2 h rocking at RT. Then sections were incubated with primary antibodies diluted in 1% BSA in PBS-Tx for 2-3 days at 4°C and washed three times in PBS-Tx prior to a 2 h incubation with fluorescence-conjugated secondary antibodies, which were diluted in 1% BSA in PBS-Tx. The stained sections were washed for 10 min with PBS-Tx, stained for 10 min with 4′,6-diamidino-2-phenylindole (DAPI; Sigma, D9542, 1:1000 in PBS), and washed extensively in PBS before mounting in RapiClear^®^ 1.52 (SUNJin Lab; RC1522001).

Whole-mount ears were fixed in 4% paraformaldehyde at room temperature for 2 h, permeabilized for 10 min with PBS-Tx, blocked with 3% BSA in PBS-Tx and then incubated for 3 h at room temperature or overnight at 4°C with primary antibodies diluted in 1% BSA in PBS-Tx. After washing, secondary antibody and DAPI staining was done as above, followed by mounting in Fluoroshield (Sigma-Aldrich, F-6182). Alternatively, DAPI staining was omitted and samples were mounted in Fluoroshield with DAPI (Sigma-Aldrich, F-6057).

The following primary antibodies were used for immunofluorescence staining: goat anti-mouse VE-cadherin (R&D Systems, AF1002; 1:200), hamster anti-mouse PDPN (Developmental Studies Hybridoma Bank, 8.1.1-a; 1:200), rabbit anti-mouse LYVE1 (Reliatech, 103-PA50AG; 1:200), rat anti-mouse EMCN (Santa Cruz Biotechnology, sc-65495; 1:200), rat anti-mouse PLVAP (BD Biosciences, 550563; 1:200), rabbit anti-mouse AQP1 (BiCell Scientific, 550563; 1:100). Secondary antibodies conjugated to Alexa Fluor 488, Alexa Fluor 594, Alexa Fluor 647, Cy3, or Cy5 were obtained from Jackson ImmunoResearch and used at a concentration of 3.75 µg/ml.

### Image acquisition

Displayed confocal images are either single or multiple tile scan images and represented as maximum intensity projections of z-stacks unless indicated differently. All confocal images were acquired using the Leica TCS SP8 or Stellaris 5 confocal microscope with the Leica LAS X software (Leica Microsystems). Following objectives were used with the Leica TCS SP8: HC FLUOTAR L 25× /0.95 W VISIR and HC PL APO CS2 63× /1.30 GLYC objectives. The Leica Stellaris 5 was used with: HC FLUOTAR L 25× /0.95 W VISIR and HC PL APO CS2 63× /1.30 GLYC objectives.

2-photon images were acquired using the Leica SP8 DIVE system with the HPX APO 20× /1.0 water dipping objective and the infrared femtosecond Ti:Sapphire (multiphoton) laser (Spectra-Physics) set to 920 nm for excitation of GFP and 1240 nm for excitation of Lectin-649.

### Scanning electron microscopy

Anesthetized animals were perfused with PBS for 2 min, followed by perfusion with EM fixative (2.5% glutaraldehyde (Sigma, G5882) and 0.1 M sodium cacodylate (Sigma, C4945) in 4% PFA (Sigma, P6148), pH adjusted to 7.4) for 2 min. The tissue was post-fixed in EM fixative for 2 h, followed by decalcification with 1:1 1 M sodium cacodylate buffer (pH 7.4) and 0.5 M EDTA (pH 8) at room temperature for five days with daily exchange of the decalcification solution. Fixed penile tissue was embedded in a gelatin-albumin matrix for vibratome sectioning. The coronal penile sections (200 µm) were kept in sodium cacodylate buffer until processing. For SEM imaging, vibratome sections were washed in 0.1M sodium cacodylate buffer (pH 7.4), post-fixed in 1 % w/v unbuffered OsO_4_ for 1 h, washed three times in water, dehydrated in EtOH (20-100%, 20% steps) and then EtOH-acetone (70:30, 50:50, 30:70, 100%; 15 min steps), critical point dried in an Agar E3000 critical point dryer (Quorum Technologies Ltd., East Essex, UK), mounted on stubs and coated with Au with an Emitech K550X sputter device (Emitech Ltd., Kent, UK) before examining using a Philips XL 30 ESEM (Philips, Eindhoven, Netherlands) operated at 15 kV accelerating voltage. Images were recorded digitally.

### Image analysis

Images were analyzed using Fiji ImageJ (version 2.0.0-rc-69/1.53c, http://imagej.nih.gov/ij/, NIH). 3D-renderings were computed from 2-photon images using the Imaris x64 software (v.9.5.1.) (Bitplane).

The sinusoidal fenestrations were characterized by measuring diameter, total area, any present gap areas with Fiji ImageJ. Fenestration frequencies were calculated using equation (1):

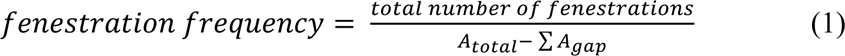

where *A_total_* is the total area of the quantified image and *A_gap_* the area of gaps presented in the quantified area, respectively. Porosity of the penile cavernous sinusoids were calculated according to equation (2):

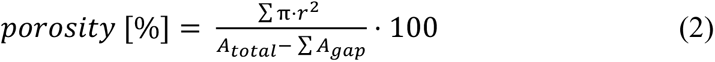

where *r* is the radius of the corresponding gap junction, A_total_ is the total area of the quantified image and A_gap_ the area of gaps presented in the quantified area, respectively. All images were acquired from one animal. Data is representative of 7 representative detail images taken from 1 section and represented as mean ± SEM.

### Flow cytometry

Mouse penises were divided into penile glans and body and cut into smaller pieces, followed by digestion in Collagenase IV (Life Technologies; 5 mg/ml), DNase I (Roche; 0.1 mg/ml) and FBS (Life Technologies; 0.5 %) in PBS at 37°C for 10-15 min, with constant shaking 950 rpm. Collagenase activity was quenched by addition of EDTA to a final concentration of 2 mM and digestion products were filtered through 50 µm nylon filters (Sysmex) and washed with FACS buffer (PBS, 0.5% FBS, 2 mM EDTA). Afterwards, cells were immediately processed for immunostaining. The Fc receptor binding was blocked with rat anti-mouse CD16/CD32 (eBioscience, 14-0161-82) followed by an incubation with antibodies (all obtained from eBioscience) against Ter119 (48-5921-82), CD45 (48-0451-82), CD11b (17-0112-82), PECAM1 (25-0311-82), and PDPN (12-5381-82). The antibody staining was followed by a FACS buffer washing step, followed by staining for dead cells using Sytox blue (Life Technologies, L34961). Cells were analyzed on a BD LSR Fortessa cell analyzer equipped with fixed laser lines (used: 405, 488, 561, and 643 nm). For compensation, the anti-rat/hamster compensation bead kit and the ArC amine reactive compensation bead kit (Life Technologies, A10628) was used. Single viable cells were gated from FSC-A/SSC-A, FSC-H/FSC-W and SSC-H/SSC-W plots and subsequent exclusion of dead cells in the violet 405 dump channel. FMO controls were used to set up presented gating schemes, allowing to distinguish and quantify distinct cell populations. Acquired flow data was processed using using FlowJo software version 10.5.0 (TreeStar).

### scRNAseq data processing

Three eight matrices of data from human penile tissue samples, containing all cell types, were obtained from the GEO database (GSE206528), with individual sample accession numbers GSM6255907, GSM6255908, GSM6255909, GSM6255910, GSM6255911, GSM6255912, GSM6255913, GSM6255914 [25]. Samples were merged and filtered to keep only cells expressing >200 genes. Mitochondrial genes and genes expressed in <3 cells were removed. In addition, cells with a fraction of mitochondrial gene counts >6% were excluded, as well as counts of long non-coding *MALAT1* RNA, which is abundantly expressed [26] and a known bias of scRNA-seq analysis. Samples were batch integrated by canonical correlation analysis (CCA) using Seurat (version 4.0.6) [27]. *Log Normalization*, dimensional reductions, graph-based clustering, UMAP visualization and differentially expressed gene (DEG) analysis were completed following Seurat v4 documentation. ECs formed separate clusters identified by their top DEGs: *EMCN*, *MMRN1*, *VWF*, *ACKR1* and *CLDN5*. EC clusters were extracted and the final dataset was obtained after cell type identification and removal of contaminants followed by re-integration and -clustering as described above. Contaminating cell types formed separated clusters of fibroblasts (*DCN*, *LUM*, *FBLN1*, *COL1A2*) or displayed interferon pathway expression patterns (*RSAD2*, *ISG15*, *IFI44L*). All analysis was performed using Rstudio (desktop version 2022.07.1) in R (version 4.2.1). A web application for data searching and visualization was generated using the shiny package of Rstudio (https://shiny.rstudio.com), and the package ShinyCell for database creation [28]. Upregulated DEGs of ED ECs compared to healthy EC were obtained for each cluster individually and further processed by GO enrichment analysis. Hypergeometric distribution test was performed using GO stats (version 2.64.0). A universal gene list was obtained from org.Hs.eg.db [version 3.16.0] and gene sets included in the analysis contained 5-1000 genes of the human genome. Pathways were considered significant when passing thresholds of P<0.05, gene count/term >10. Additionally, relevant GO terms were selected by OddsRatios and classified in seven different annotation categories, visualized using ggplot2 (version 3.4.2). Trajectory analysis was performed (curve not shown) using SCORPIUS (version 1.0.8) [29], using default parameters and *k =* 5. Cells were mapped along the trajectory. Cell order along the trajectory was retained when visualizing selected genes in a heatmap.

### Data presentation and statistical analysis

GraphPad Prism 9 was used for data visualization and statistical testing. Data between two groups were compared using the unpaired t-test with Welch’s correction. Differences were considered statistically significant when *p*<0.05 and indicated on the graphs with star symbols: * *p* < 0.05, ** *p* < 0.01 and *** *p* < 0.001. Hypergeometric distribution was assessed for GO enrichment analysis. DEG analysis was performed using Wilcoxon rank sum test.

## Results

### Transcriptomics of human penile cavernous vasculature identifies EC populations with hybrid vessel identity and distinct phenotypic change in erectile dysfunction

To characterize the molecular features of penile cavernous (pc) vasculature, we extracted endothelial cell (EC) transcriptomes, identified based on the expression of pan-endothelial markers (*PECAM1*, *CDH5*), from single cell RNA sequencing (scRNA-seq) data obtained from human corpus cavernosum tissue [25]. The dataset included samples from men without erectile dysfunction (n=3, healthy), as well as from non-diabetic (n=3, ED_nDM) and type 1 diabetic (n=2, ED_DM) men treated for erectile dysfunction (ED) (**Fig. 1A**).

**Figure 1.**
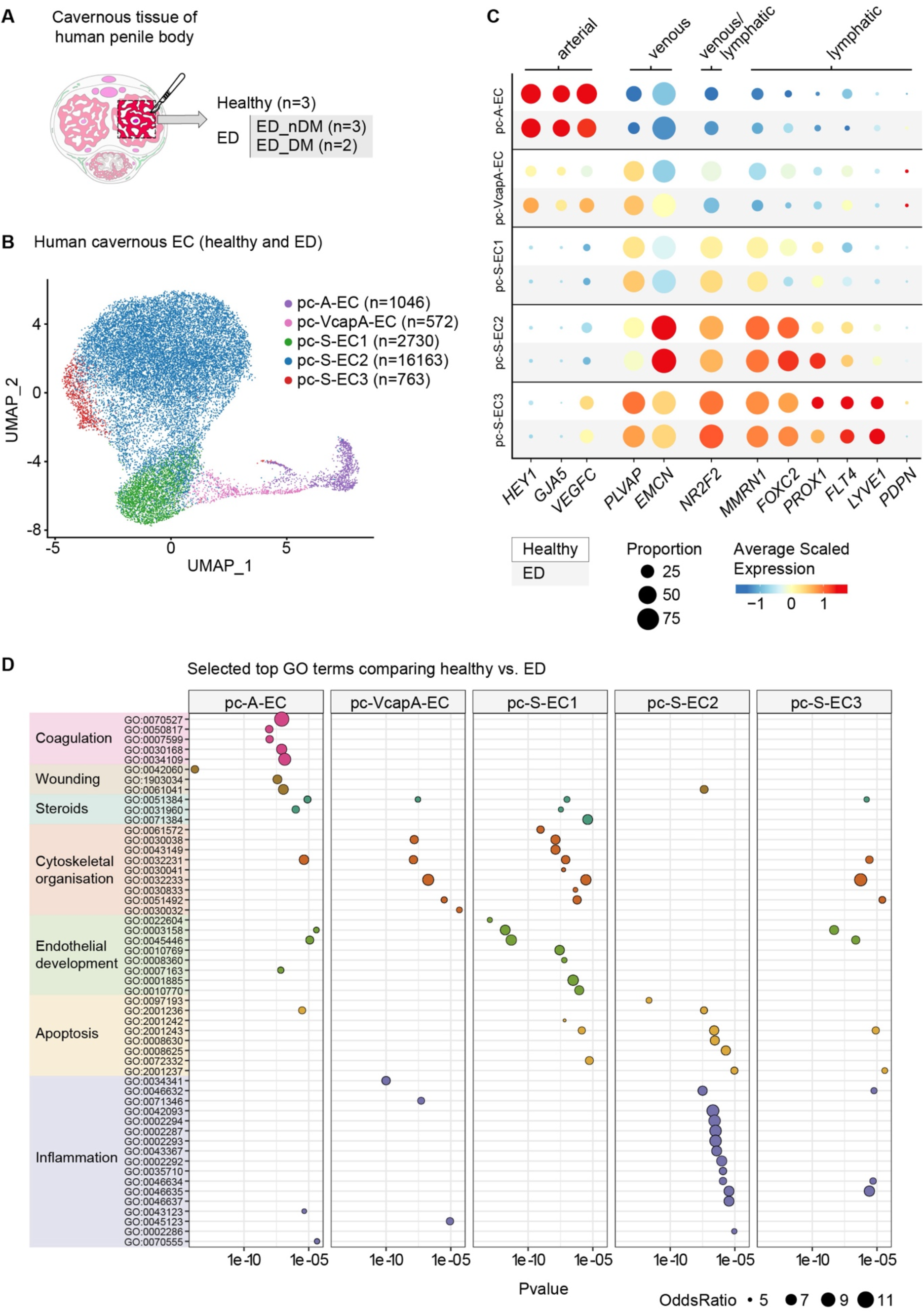
Single cell transcriptomics of human penile cavernous ECs in health and in erectile dysfunction. (**A**) Schematic presentation of human penile cavernous tissue used for single cell RNA sequencing analysis. Analyzed dataset [25] included samples from men without erectile dysfunction (ED) (n=3, healthy), as well as from non-diabetic (n=3, ED_nDM) and type 1 diabetic (n=2, ED_DM) men treated for erectile dysfunction. (**B**) The five penile cavernous EC (pc-EC) clusters, from healthy and ED tissue samples, visualized in a UMAP landscape and colored by cluster assignment. (**C**) Dot plots showing selected EC subtype markers for the five EC clusters presented in (B) between healthy and ED pc-ECs. Dot size illustrates percentage of cells presenting transcript sequence counts and color illustrates the average scaled gene expression level (log2 fold change) within a cluster. (**D**) GO analysis of genes enriched in the clusters of the ED pc-ECs. Dot size illustrates OddsRatio and color illustrates superordinate categories, with corresponding terms ordered by P value along the X-axis.

After removal of contaminating cells identified as fibroblasts and mural cells, which are commonly found in scRNA-seq datasets of vascular cells [30,31], we obtained in total 21 274 high-quality ECs. They distributed into five clusters after applying Harmony method for batch correction and Seurat graph-based clustering approach (**Fig. 1B**). ECs from individual samples and health states contributed to each cluster (**Fig. S1**). Based on known molecular signatures and in line with previous analysis [25,32], we annotated one cluster of penile cavernous arterial ECs (pc-A-ECs) expressing arterial markers (e.g. *HEY1*, *GJA5*, *VEGFC*), and three clusters of penile cavernous sinusoidal ECs (pc-S-EC1-3) expressing venous markers (e.g. *EMCN*, *PLVAP, NR2F2*) (**Fig. 1B and C**). An additional cluster was characterized by heterogenous expression of EC identity markers, potentially representing transition along venous-capillary-arterial trajectory (pc-VcapA-EC) (**Fig. 1B and C**). Ordering of cells based on similarities in their gene expression patterns generated a linear trajectory that indeed placed pc-VcapA-EC cluster in between pc-S-EC1 and pc-A-EC clusters, and revealed zonated expression of arterial, capillary and venous markers previously identified in the brain vasculature [30] (**Fig. S1)**. Unexpectedly, we found that the pc-S-EC clusters also expressed varying degrees of *PROX1,* which is the master regulator of LEC fate, along with other markers of lymphatic and hybrid EC identity (e.g. *FOXC2*, *FLT4, LYVE1*) (**Fig. 1C**). However, The data will be made available for querying upon publishing.

We observed no major differences in the expression of venous and arterial EC identity markers in ED compared to healthy tissue (**Fig. 1C**). In contrast, the two clusters with hybrid EC identity, pc-S-EC2 and pc-S-EC3, displayed an increased expression of the lymphatic markers *PROX1,* as well as *FLT4* and *LYVE1*, respectively, in ED compared to healthy state (**Fig. 1C**). To investigate pathological gene expression changes in pc-ECs globally, we performed Gene Ontology (GO) analysis of the upregulated, differential expressed genes (DEGs) between ED and healthy pc-EC clusters. This analysis revealed a distinct disease-associated profile in the different EC types, with selective enrichment of biological processes related to coagulation in pc-A-EC cluster, and inflammation in pc-S-EC2 cluster (**Fig. 1D**). Other pc-EC clusters shared a more similar response, characterized by enrichment of processed related to steroid response, cytoskeletal organization and endothelial development (**Fig. 1D**).

Taken together, these results reveal zonation of human corpus cavernosum vasculature, and identify cavernous penile sinusoidal endothelium as a new hybrid vascular bed with a distinct erectile dysfunction-associated change in gene expression.

### Murine penile cavernous sinusoids are blood-perfused hybrid vessels

To investigate if the molecular features of penile cavernous sinusoids (pc-Ss) are conserved in the structurally similar murine penile vasculature, we analyzed the expression of EC identity markers using genetic reporter mice and immunofluorescence staining in 60-90 µm coronal sections of the penile glans and body. The cavernous tissues analyzed included corpus cavernosum glandis (CCg) [33], the bilateral MUMP corpora cavernosa [14] and bilateral corpora cavernosa uretrae (CCug) in the penile glans, as well as the corpora cavernosa uretrae (CCu) and the bilateral corpus cavernosa (CC) in the penile body [15] (**Fig. 2A**).

**Figure 2.**
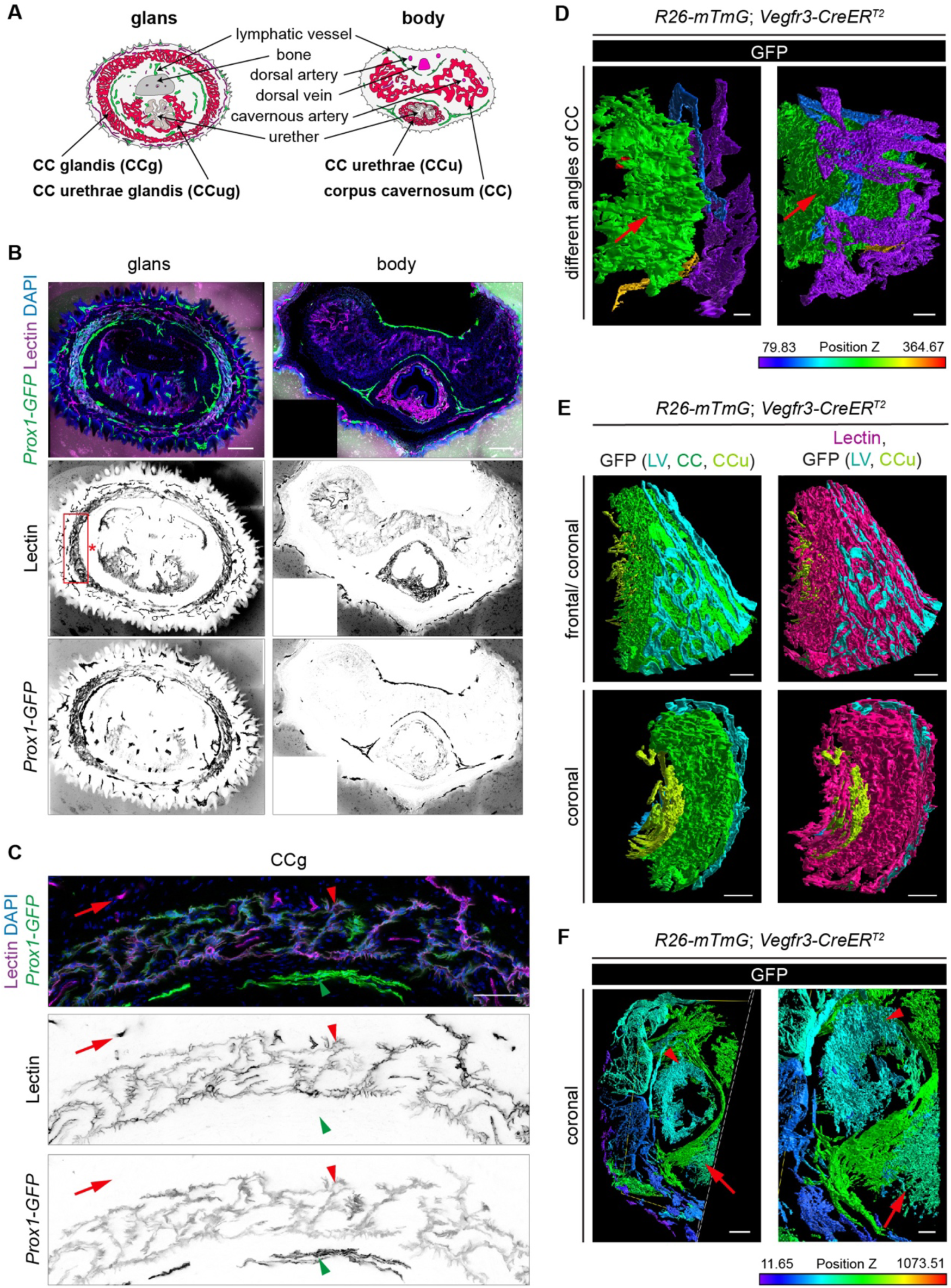
Characterization of the murine penile vasculature. (**A**) Schematic presentation of the murine penile anatomy. Blood vessels (pink), penile cavernous sinusoids (pc-Ss) (red), lymphatic vessels (green), baculum (penis bone, gray) and ureter (brown) are displayed. CCg, corpus cavernosum glandis; CCug, corpus cavernosum urethrae glandis; CCu, corpus cavernosum urethrae; CC, corpus cavernosum. (**B, C**) Vascular blood perfusion (visualized by lectin perfusion) and expression of *Prox1-GFP* in the penile glans and body. A magnified view of the boxed region in (B) is shown in (C). Red arrowheads point to blood perfused *Prox1-GFP* positive sinusoids, and blood vessels (red arrows) and lymphatic vessels (green arrowheads) are indicated. (**D-F**) Imaris rendering from 2-Photon images of cavernous tissue from tamoxifen treated *R26-mTmG;Vegfr3-CreER^T2^*mice, displayed in colored depth-coding and showing different angles of a coronal section of CC (arrow) (D). Note colocalization of GFP (*Vegfr3* expression) with intravenously injected lectin (purple) in CC and CCu, but not in lymphatic vessels (LV) (E). (F) CCu (arrowhead) and CC (arrow) are displayed. Scale bars, 50 μm (**C**), 80 μm (**D**), 100 μm (**F** (right panel)), 200 μm (**B, E, F** (left panel)).

The blood and lymphatic vessels were identified via intravenous injection of biotinylated Lectin, in combination with the visualization of the LEC marker *Prox1* using the *Prox1-GFP* transgene (**Fig. 2B**). In addition to the GFP^+^ Lectin^-^ lymphatic vessels and GFP^-^Lectin^+^ blood vessels, we observed Lectin^+^ i.e. blood perfused pc-Ss that unexpectedly also expressed *Prox1-GFP* (**Fig. 2B and C**). Pc-Ss expressed another key LEC marker *Vegfr3*, as detected by GFP expression in a tamoxifen-treated *R26-mTmG;Vegfr3-CreER^T2^* reporter mouse (**Fig. 2D-F**). Three dimensional (3D)-rendering of 2-photon z-stack reconstruction of GFP^+^ penile vasculature in *R26-mTmG;Vegfr3-CreER^T2^* mice revealed CC (**Fig. 2D and E**) and CCu (**Fig. 2F**), positioned ventral to the lumen and extending from distally into the urethral flaps. Intravenously injected Lectin was found in *Vegfr3* expressing blood perfused pc-Ss, but not in lymphatic vessels (**Fig. 2E**). These results show that murine pc-Ss are blood-perfused vessels expressing the key LEC signature genes *Prox1* and *Vegfr3*, suggesting conservation of their hybrid identity between mouse and man.

### Heterogeneity, acquisition and maintenance of penile sinusoid identity

All murine penile cavernous tissues analyzed expressed the expected pan-endothelial markers, including junctional proteins *Cdh5* (encoding VE-cadherin) (**Fig S2**, **Table 1**), as well as the venous EC marker EMCN (**Fig. 3A, Fig. S2**). While the majority of pc-S-ECs in CCug and CCg of the penile glans showed prominent *Prox1-GFP* fluorescence (**Fig. 3A**), a smaller fraction of pc-S-ECs in the CCu and CC of the penile body were GFP^+^ (**Fig. S2**). We also observed heterogenous expression of the lymphatic vessel endothelial hyaluronan receptor 1 (LYVE1), while PDPN, the bona fide marker of mature lymphatic vessels, was not expressed in pc-S-ECs (**Fig. 3B**, **Fig. S2**, **Table 1**).

**Figure 3.**
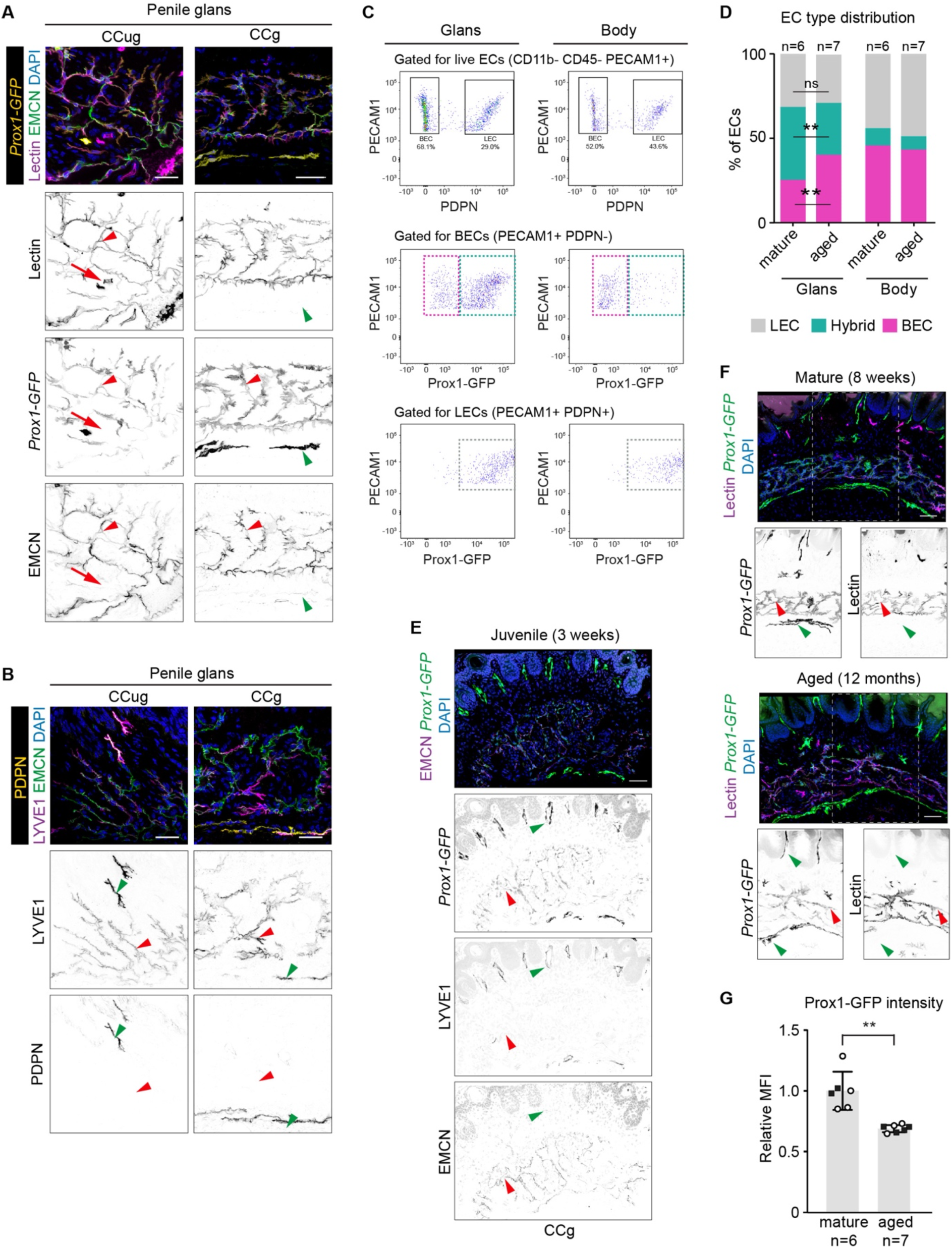
*Prox1* expression in penile cavernous sinusoids of juvenile and aged mice. **(A)** *Prox1-GFP*expression, blood perfusion (determined by lectin perfusion) and immunofluorescence for EMCN is displayed in transversal sections of the penile glans (CCug and CCg). Veins (red arrows), lymphatic vessels (green arrowheads) and pc-Ss (red arrowheads) are indicated. (**B**) Immunofluorescence for LYVE1 and PDPN in vibratome sections of the glans (CCug and CCg). Lymphatic vessels (green arrowheads) and pc-Ss (red arrowheads) are indicated. (**C, D**) Representative gating schemes (C) and summary graph (D) for the analysis of BECs, LECs and *Prox1^+^* hybrid ECs in the penile glans (left panels) and body (right panels) of mature and aged mice. Data is representative of n=6 (mature) and n=7 (aged) mice from 2 independent experiments. (**E**) *Prox1-GFP* expression and immunofluorescence for LYVE1 and EMCN in CCg of juvenile (P21) mice. (**F**) *Prox1-GFP* expression and blood perfusion (determined by lectin perfusion) in CCg of mature (8-week-old) and aged (12-month-old) mice. Note expression of *Prox1* in pc-Ss at all stages (red arrowheads), as well as in lymphatic vessels (green arrowheads). (**G**) Quantification of median fluorescent intensity (MFI) of *Prox1-GFP* expression in hybrid ECs (PECAM1^+^ PDPN^-^ *Prox1-GFP*^+^) of the glans of mature (8-10-week-old) and aged (10-12-month-old) mice by flow cytometry. Data represents mean (n=6 (mature) and n=7 (aged) mice from 2 independent experiments, indicated by different symbols) ± SEM. Unpaired t-test with Welch’s correction, \*\**P* < 0.01. Scale bars, 25 μm (**A, B**), 50 μm (**E**, **F**).

**Table 1.** Endothelial cell marker expression in penile cavernous sinusoids. The analyzed markers in ECs of penile cavernous sinusoids (pc-Ss) in comparison to previously reported expression in hybrid vessels: Schlemm’s canal (SC) [2–4,34–39], ascending vasa recta (AVR) [5,40–42] and remodeled spiral artery (rSA) [6,43–46]. (+)=expression detected, (−)=not detected, (+/-)=discontinuous expression, nrf=no reporting found.

| Specificity | Marker | pc-Ss |  |  |  | SC | AVR | rSA |
| --- | --- | --- | --- | --- | --- | --- | --- | --- |
|  |  | CCug | CCg | CCu | CC |  |  |  |
| Pan-EC | PECAM1 | + | + | + | + | + | + | + |
|  | TIE2 | + | + | + | + | + | + | + |
|  | VE-cadherin | + | + | + | + | + | nrf | + |
|  | VEGFR2 | + | + | + | + | + | - | nrf |
| BEC | AQP1 | + | + | + | + | + | - | + |
|  | CD34 | + | + | + | + | + | + | nrf |
|  | PLVAP | + | + | + | + | + | + | nrf |
|  | EMCN | + | + | + | + | + | + | - |
|  | vWF | + | + | + | + | + | nrf | + |
| LEC, hybrid | PROX1 | + | + | +/- | +/- | + | + | + |
|  | VEGFR3 | + | + | + | + | + | + | + |
|  | LYVE1 | + | + | + | + | - | - | ~ |
| LEC | CCL21 | - | - | - | - | + | nrf | - |
|  | PDPN | - | - | - | - | - | - | - |

Flow cytometry analysis of the murine penile tissue confirmed the presence of three PECAM1^+^ EC populations with blood (*Prox1-GFP^-^*PDPN^-^), lymphatic (*Prox1-GFP^+^*PDPN^+^) and hybrid (*Prox1-GFP^+^*PDPN^-^) vessel identities (**Fig. 3C**). Interestingly, we observed a relative decrease in the *Prox1^+^* hybrid EC population (p=0.0031, unpaired Student’s t-test with Welch correction), with a corresponding increase in the relative *Prox1^-^* BEC population (p=0.0024) in penile glans of aged (10-12 months of age, n=7) in comparison to mature (2-3 months of age, n=6) mice (**Fig. 3D**).

The murine penis develops postnatally, with well-defined cavernous tissues being distinguishable from postnatal day (P)10 in the penile glans and body [33]. Analysis of penile vasculature of juvenile 3-week-old mice showed EMCN^+^ pc-Ss that were weakly positive for *Prox1-GFP* fluorescence, but did not yet express LYVE1 (**Fig. 3E**). To study the maintenance of pc-S-EC identity during ageing, we analyzed vessel architecture and *Prox1* expression in the penile tissues of mature (2-3 months) and aged (10-12 months) mice. Immunofluorescence analysis did not reveal differences in the morphology or blood perfusion of pc-Ss between the developmental stages, but *Prox1-GFP* fluorescence appeared weaker in aged mice (**Fig. 3F**). This finding was confirmed by flow cytometry analysis of hybrid ECs showing lower median fluorescence intensity (MFI) of *Prox1-GFP* in penile glans of aged in comparison to young mice (**Fig. 3G**). Interestingly, decline in PROX1 expression was previously associated with age-related deterioration of Schlemm’s canal, the hybrid vessel of the eye [3].

To assess conservation of pc-S-EC heterogeneity in man, we processed transcriptomes of ECs from healthy human corpus cavernosum tissue that distributed into five clusters, representing aortic and sinusoidal ECs (**Fig. S3**). As expected, all clusters expressed the pan-EC markers *PECAM1* and *CDH5*, but showed variable expression of the tight-junction molecule *CLDN5* (**Fig. S3**). Additional cluster markers indicated further molecular heterogeneity within the pc-S-EC population, in line with the previous studies [25,32], including high expression of the water channel aquaporin 1 (AQP1) in the pc-S-EC1 cluster (**Fig. S3**). A similar heterogeneous expression of *Cldn5* and AQP1 (**Fig. S3**) within cavernous endothelium was also observed *in vivo* in murine penile tissue.

Collectively, these results show that mouse and human pc-Ss are molecularly heterogenous and exhibit an identity similar to that of the previously described hybrid vasculatures of the eye [2–4], renal medulla [5], and placental decidua [6]. *Prox1* expression in pc-Ss is initiated prior to sexual maturity, and largely maintained throughout life, though decreasing with age.

### Ultrastructural and functional features of murine penile sinusoids

Pc-Ss control the rigidity of the penis by entrapping incoming arterial blood, engorging and containing the blood within the sinusoidal spaces, which exerts pressure on the tunica albuginea and the occlusion of traversing veins. The sinusoidal endothelium is thus exposed to a range of pressure, stretch and shear forces likely to impact on their cell-cell junctions. Staining of whole-mounted tissue revealed continuous VE-cadherin positive cell-cell junctions along torturous cell borders in LYVE1 positive pc-S-ECs in CCu (**Fig. 4A**) and CC (**Fig. S4**). To characterize the ultrastructural features of pc-Ss, we performed scanning electron microscopy of vibratome sections (**Fig. 4B**). Analysis of the luminal surface of pc-S-ECs revealed interdigitating and overlapping cell-cell contacts and protrusions, as well as numerous intercellular gaps (**Fig. 4C**). Consistent with the sinusoidal nature of pc-Ss, we observed abundant fenestrations (0.17 fenestrations/µm^2^ with 0.15% porosity and a diameter of 40 – 400 nm (mean 92.15 ± 2.89 nm, n=319)) (**Fig. 4D**) within the CC of the penile body. Expression of the plasmalemma vesicle-associated protein (PLVAP) in pc-S-ECs of all cavernous tissues analyzed suggested the presence of filter-like diaphragms, previously found in fenestrae and trans-endothelial channels that confer vascular barrier function (**Fig. S4**). However, PLVAP is also expressed on ECs without diaphragms, such as liver sinusoidal ECs that maintain transcellular pores of similar diameters (50 – 250 nm, mean 90.7 ± 11.7 nm), but higher density (8.45 fenestrations/µm^2^) and porosity (5.93%) [47,48] compared to pc-S-ECs.

**Figure 4.**
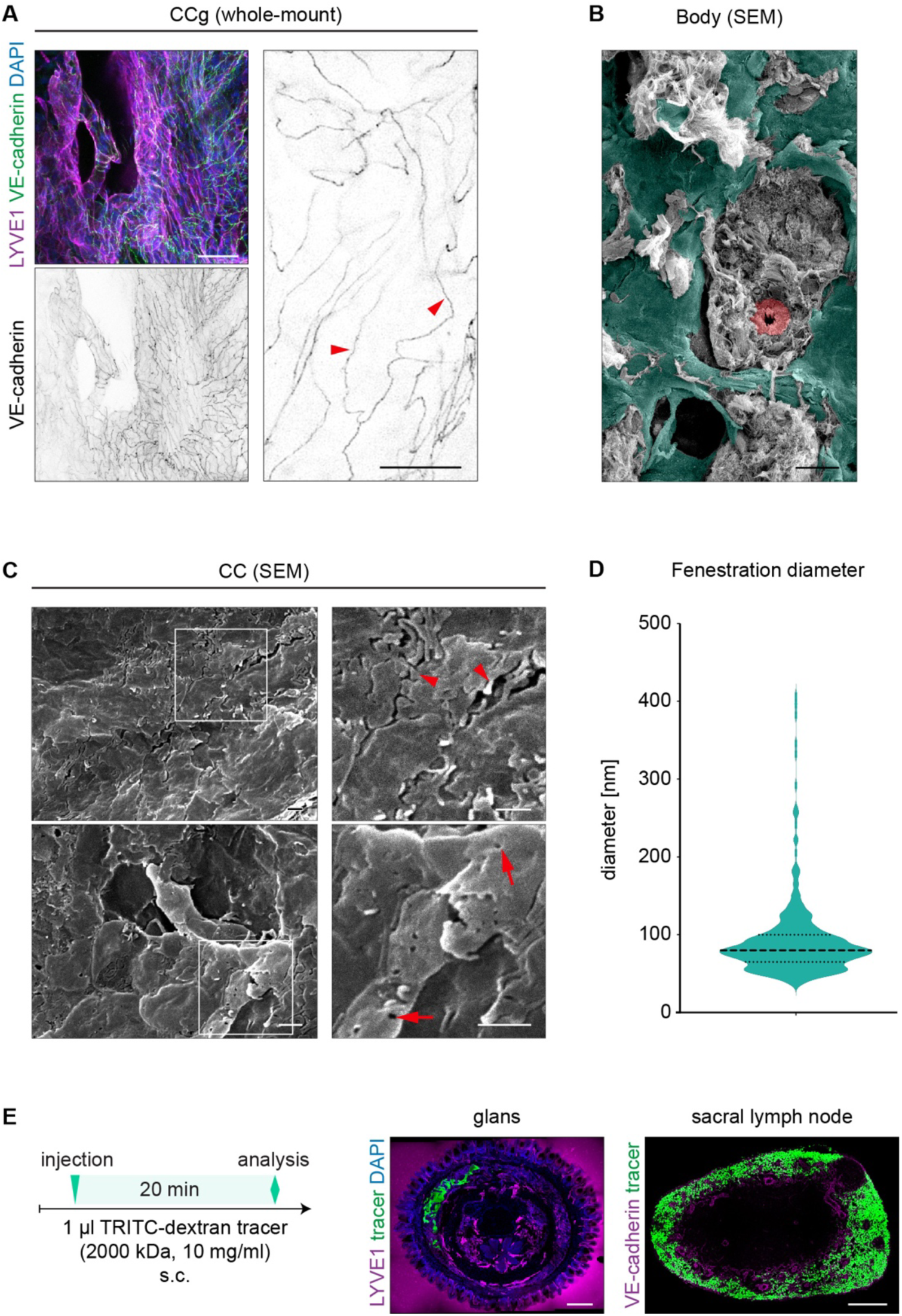
Ultrastructural and functional features of murine penile cavernous sinusoids. (**A**) Whole-mount immunofluorescence of CCu. Note the presence of continuous VE-cadherin^+^ cell-cell junctions (red arrowheads). (**B, C**) SEM analysis of the luminal surface of sinusoidal endothelium (color-coded in cyan in B), showing interdigitating and overlapping cell-cell contacts and protrusion (C, red arrowheads), and intercellular gaps and fenestrations (C, red arrows). (**D**) Quantification of fenestration size in pc-Ss of CC. Data is representative of 7 images taken from 1 section and represented as mean ± SEM. (**E**) Experimental plan for the injection of TRITC-dextran tracer (2×10^6^ MW; left panel), uptake of the tracer by the pc-Ss in the glans (middle panel) and transport of the tracer to the sacral lymph node (right panel) is displayed. Scale bars, 1 μm (**C**), 25 μm (**A** (right panel), **B**), 50 μm (**A,** left panel), 250 μm (**E**).

To functionally assess potential macromolecular uptake by pc-Ss, we injected high-molecular weight dextran tracer (2×10^6^ MW) subcutaneously into the penile glans. As expected [49,50], the tracer was drained by lymphatic vessels to lumbar and sacral lymph nodes (**Fig. 4E**). While the tracer is too large to enter blood capillaries, it was unexpectedly detected within pc-Ss (**Fig. 4E**). The ultrastructural and functional analyses thus indicate a potential role of penile sinusoids in fluid and macromolecule uptake, similar to that of lymphatic vessels.

### VEGF-C is not required for development of the penile hybrid vasculature in mice

The development of all lymphatic and previously described hybrid vascular beds is dependent on vascular endothelial growth factor C (VEGF-C) signaling via its receptor VEGFR3 [2,6]. To test if the regulatory mechanisms are conserved in pc-S formation, we inhibited VEGF-C using adeno-associated vectors (AAVs) encoding a soluble ligand binding domain of VEGFR3 fused to the IgG Fc domain (mVEGFR3_1-4_-mFc) that acts as a ligand trap (VEGF-C trap) [51]. AAVs were administered to P6 pups, prior to the development of the cavernous tissues, by intraperitoneal injection and the penile tissue was analyzed at 5 weeks of age (**Fig. 5A**). Control mice were treated with AAVs encoding the non-ligand binding region of the VEGFR3 extracellular domain (mVEGFR3_4-7_-mFc). In accordance with previous studies [24], this led to a systemic production of the trap molecule (**Fig. S5**) and blocked VEGF-C-induced lymphangiogenesis in the ear (**Fig. S5**). We also observed inhibition of the formation of penile lymphatic vessels (**Fig. 5B**), which was associated with tissue swelling (**Fig. S5**). VEGF-C trap-treated mice additionally displayed developmental defects of the penis (**Fig. S5**) and phenotypic characteristics of cryptorchidism, with testicles being positioned suprascrotally inside the abdomen (**Fig. S5**). However, no apparent differences in the morphology of pc-Ss or the expression of pc-S-EC markers were observed in mice treated with AAVs encoding the VEGF-C trap compared to control AAVs (**Fig. 5C**). This demonstrates that the development of penile cavernous sinusoids and acquisition of their identity is independent of VEGF-C signaling.

**Figure 5.**
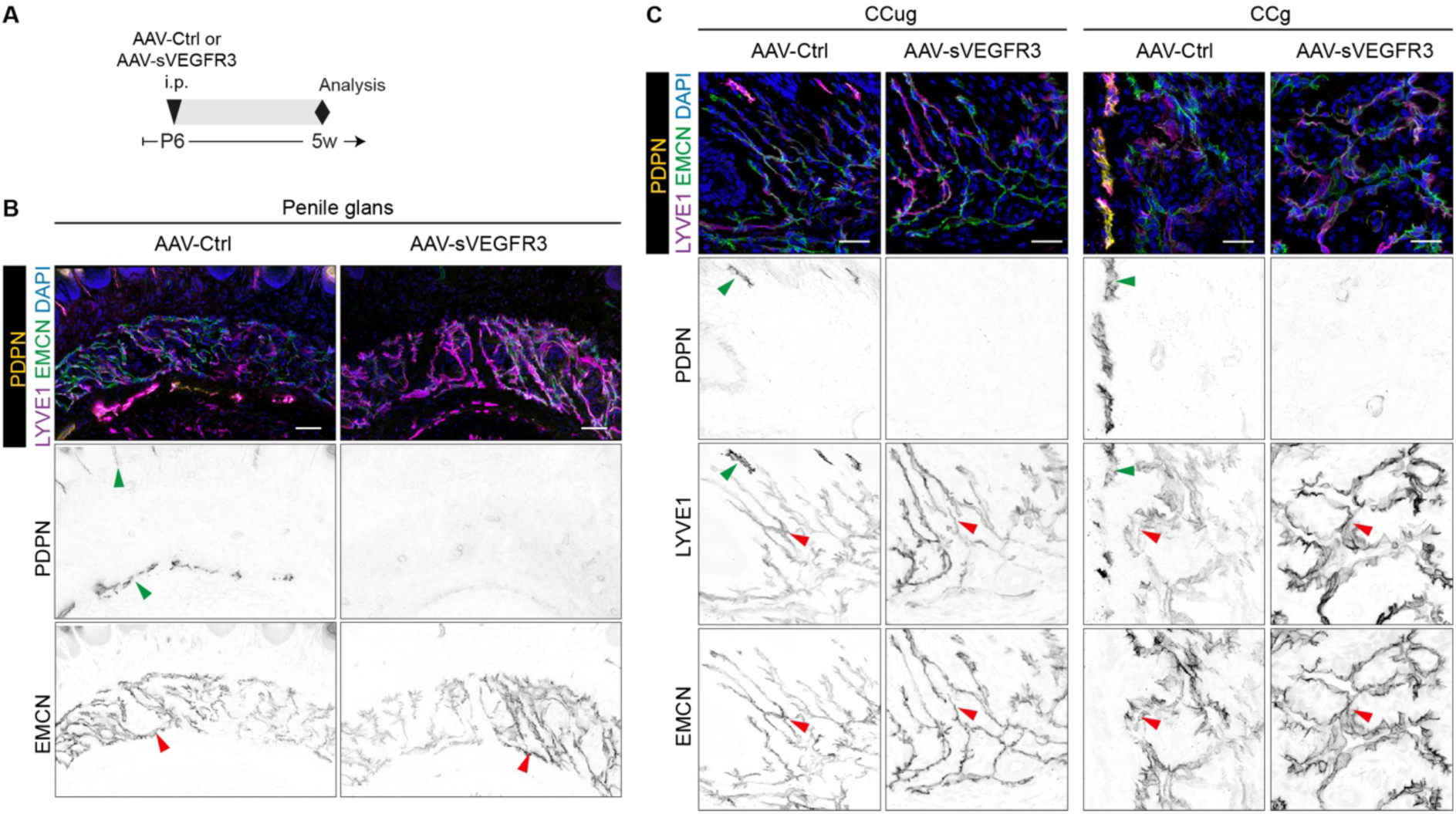
The lack of effect of VEGF-C inhibition on the development of murine penile cavernous sinusoids. (**A**) Treatment scheme for inhibition of VEGF-C signaling prior to cavernous tissue development using the soluble VEGF-C trap (AAV-sVEGFR3) or control trap (AAV-Ctrl). (**B, C**) Immunofluorescence of vibratome sections of penile glans showing absence of PDPN^+^ lymphatic vessels (green arrowheads) but normal appearance of EMCN^+^LYVE1^+^ pc-Ss (red arrowheads) in AAV-sVEGFR3-treated animals. Scale bars, 25 μm (**C**), 50 μm (**B**).

## Discussion

ECs of blood and lymphatic vessels represent differentiated cell lineages defined by expression of unique identity markers. Recently, hybrid vasculatures that possess features and functions of both blood and lymphatic vessels have been identified in multiple tissues, including the Schlemm’s canal in the eye. This has enabled identification of new molecular targets for the treatment of glaucoma [52–55], which is associated with defective Schlemm’s canal function [56–58]. In this study, we identify the penile cavernous sinusoids as a new hybrid vasculature that is characterized by the expression of key LEC regulators, including *Prox1*, and show distinct phenotypic alterations in ED.

Considering that ED is predominantly a disease of vascular origin, there is a striking lack of understanding of the specialized penile vasculature. In the non-erected, flaccid state the pc-Ss are minimally blood perfused. During an erection, pc-Ss fill, engorge and retain blood, thereby facilitating the tissue rigidity required for successful mating. This uniquely exposes pc-Ss to highly varying interstitial pressure, stretching and shear stress, which is likely to impact their gene expression. Previous transcriptome studies of ECs of human penile cavernous tissue [30,31] identified populations of arterial and venous/sinusoidal ECs, with the latter showing high heterogeneity as also observed by us. Unexpectedly, we additionally found that human and murine pc-S-ECs express the master regulators of LEC fate and lymphangiogenesis, *Prox1* and *Vegfr3*, respectively. Moreover, similar to the previously identified hybrid vasculatures [1], they did not express the marker of mature lymphatic vessels, PDPN. We further found that murine pc-Ss acquire hybrid vascular identity prior to sexual maturity, but the expression of *Prox1* in pc-S-ECs of the penile glans declined in aged in comparison to young mice. Notably, an age-related and pathological decline in PROX1 expression was described in the Schlemm’s canal and correlated with its functionality [3]. Tight control of PROX1 levels is also required for the maintenance of LEC identity and lymphatic vessel function [59]. Hence, it will be of interest to understand if regulation of hybrid identity of pc-S-ECs correlates with the functionality of pc-Ss and erectile function during aging.

Both murine and human CCu and CC are devoid of a lymphatic vascular system responsible for the drainage of excess interstitial fluid in most other tissues. We found that pc-Ss can likely compensate for this function by their ability to take up high-molecular weight tracers, which is consistent with the presence of fenestrae and AQP1 water channels in pc-S-ECs. Interestingly, vascular access and resuscitation via the corpus cavernosa has been established as an effective route in hypovolemic males [60,61], but no further investigations have been conducted on the mechanisms of saline uptake by pc-Ss.

Unlike other lymphatic and hybrid vasculatures analyzed so far [2,6,42], we found that the development of the murine penile hybrid vasculature was not dependent on VEGF-C signaling. Neonatal inhibition of VEGF-C using AAV-encoded soluble VEGFR3, concomitant with the inhibition of penile lymphatic vessel formation, unexpectedly resulted in defective penile development and phenotypic characteristics of cryptorchidism. While these unexpected phenotypes are interesting, the underlying mechanisms remain unclear. Interestingly, cases of cryptorchidism were described in patients with Noonan syndrome, lymphedema distichiasis and Hennekam syndrome [62,63], suggesting a potential link between lymphatic and testicular development. It is also possible that VEGF-C targets nonendothelial cells in the male genital tract, as was previously reported for VEGF that is able to bind its receptors VEGFR1 and VEGFR2 in the Leydig cells of the testis [64]. Studies on the mechanisms of pc-S development and cellular origin of pc-S-ECs using inducible Cre/loxP-based approaches are hampered due to sensitivity of penile development to the anti-estrogen tamoxifen used as an inducing agent [16,65]. Tamoxifen treatment during neonatal period leads to defects in the formation of epithelial spines of the penile glans and the bone, and cavernous tissue atrophia ([66]; and unpublished data).

In summary, our study establishes pc-Ss as a new specialized hybrid vasculature that exhibits distinct phenotypic alterations in patients with erectile dysfunction, characterized by an upregulation of lymphatic markers and altered expression of genes related to immune regulation. Our characterization of pc-Ss and the associated searchable Web application for exploring transcriptome data of pc-ECs may help identify new molecular targets for the treatment of conditions associated with dysregulated penile vasculature, such as ED.

## Acknowledgements

We thank Geoffrey Daniel (Swedish University of Agricultural Sciences, Uppsala) for SEM imaging and support in sample preparation. We also thank Sagrario Ortega (CNIO, Madrid) for the *Vegfr3*-*CreER^T2^*mice, Ralf Adams (Max Planck Institute for Molecular Biomedicine, Münster) for the *Cdh5*-*CreER^T2^* mice, Christer Betsholtz (Uppsala University, Uppsala) for the *Cldn5-GFP* mice, and Kari Alitalo (University of Helsinki, Helsinki) for the AAVs. BioVis facility (Uppsala University) is acknowledged for flow cytometer and microscopy usage and support, and Sofie Lunell Segerqvist and Aissatu Mami Camara for technical assistance.

