## Supplementary Figures 1-5 for "Penile cavernous sinusoids are Prox1-positive hybrid vessels"

**Figure S1. Characterization of human pc-EC clusters in scRNA-seq datasets of healthy and ED cavernous tissue.** (**A**) The distribution of pc-ECs from healthy and ED cavernous tissues in a UMAP landscape, showing contribution of cells from the different health states to all clusters. (**B**) Heatmap showing zonation of arterial, capillary and venous markers across the pc-A-EC, pc-VcapA-EC and pc-S-EC1 clusters. Each column represents one cell ordered along a trajectory drawn through the pc-A-EC, pc-VcapA-EC and pc-S-EC1 clusters (SCORPIUS). Red color indicates high, blue color low ex pression. (**C**) Violin plots showing the expression (Y-axis) of lymphatic markers PROX1, FLT4 and LYVE1 in the five pc-EC clusters (X-axis) from healthy (blue) and ED (red) cavernous tissue. Dots represent individual cells, and shape shows their distribution.

**Figure S2. Expression of vascular endothelial markers in penile cavernous sinusoids.** (**A**) Analysis of Cdh5 (encoding VE-cadherin) expression using Cdh5-CreERT2 mice in combination with the R26-tdTom reporter. Genetic constructs and scheme for 4-hydroxytamoxifen are shown. Note colocalization of tdTom (i.e. Cdh5) expression with intravenously injected lectin (indicating blood perfusion) in pc-Ss (red arrowheads). (**B**) Prox1-GFP expression, blood perfusion (determined by lectin perfusion) and immunofluorescence for EMCN is displayed in transversal sections of the penile body (CCu and CC). Veins (red arrows) and pc-Ss (red arrowheads) are indicated. (**C**) lmmunofluorescence for LYVE1 and PDPN in vibratome sections of the penile body (CCu and CC). Lymphatic vessels (green arrowheads) and pc-Ss (red arrowheads) are indicated. Scale bars, 25 µm (**B**, **C**), 200 µm (**A**).

**Figure S3. Molecular features of penile cavernous sinusoids.** (**A**) ECs of five healthy, human cavernous EC clusters visualized in a UMAP landscape and colored by cluster assignment. (**B**) Dot plots of selected EC subtype markers for the five EC clusters presented in (A). (**C**) Violin plots showing the distribution of gene expression level (Y-axis) of selected pan-EC marker genes (PECAM1, CLDN5, CDH5) and mural cell marker gene (PDGFRB) in healthy penile cavernous EC subtypes (X-axis). (**D**) Dot plots of selected differentially expressed genes presented for the four pc S-EC clusters 1-4 presented in (A). Dot size illustrates percentage of cells presenting transcript sequence counts and color illustrates the average scaled expression level (log2 fold change) within a cluster. (**E**) Cldn5-GFP expression and colocalization with intravenously injected lectin (indicating blood perfusion) in the pc-Ss (red arrowheads) of the glans (CCug and CCg, left panel) and body (CCu and CC, right panel). Green arrowhead points to a non-perfused Cldn5-GFP+ lymphatic vessel. (**F**) Immunofluorescence of vibratome sections of the penile glans for AQP1. Higher magnification images of boxed regions with AQP1+ ECs (red arrowheads) in F are shown on the right. Scale bars, 5 µm (**F** (details), 25 µm (**E**, **F** (overviews)).

**Figure S4. Molecular features of penile cavernous sinusoids.** (**A**) Whole-mount immunofluorescence of CC in the penile body. Note the presence of continuous VE-cadherin+ cell-cell junctions (red arrowheads). (**B**) Immunofluorescence of vibratome sections of the penile glans for PLVAP. Scale bars, 25 µm (**A** (right panel), **B**), 100 µm (**A**, left panel).

**Figure S5. The effect of VEGF-C inhibition on genital development.** (**A**) Western blot analysis showing soluble AAV-sVEGFR3 (VEGFR3$_{1\text{-}4}$-Ig) or the AAV-Ctrl (VEGFR3$_{4\text{-}7}$-Ig) in serum four weeks after intraperitoneal injection of AAV vectors. Data are representative of n=4 (AAV-sVEGFR3), n= 5 (AAV ctrl) and n= 1 (untreated) mice from 3 independent experiments. (**B**) Whole-mount immunofluorescence of ears showing inhibition of lymphatic vessel forma-tion in AAV-sVEGFR3-treated mice. (**C-E**) Morphological assessment of genital development after AAV treatment. Note the development of edematous swelling of the penile glans (C), failure to extend out of the external prepuces (D), and cryptorchidism (E) in the VEGF-C trap (AAV-sVEGFR3) treated penile glans. Scale bars, 500 µm (**B**), 1 mm (**C**, **D**), 5 mm (**E**).

**S1 Fig**

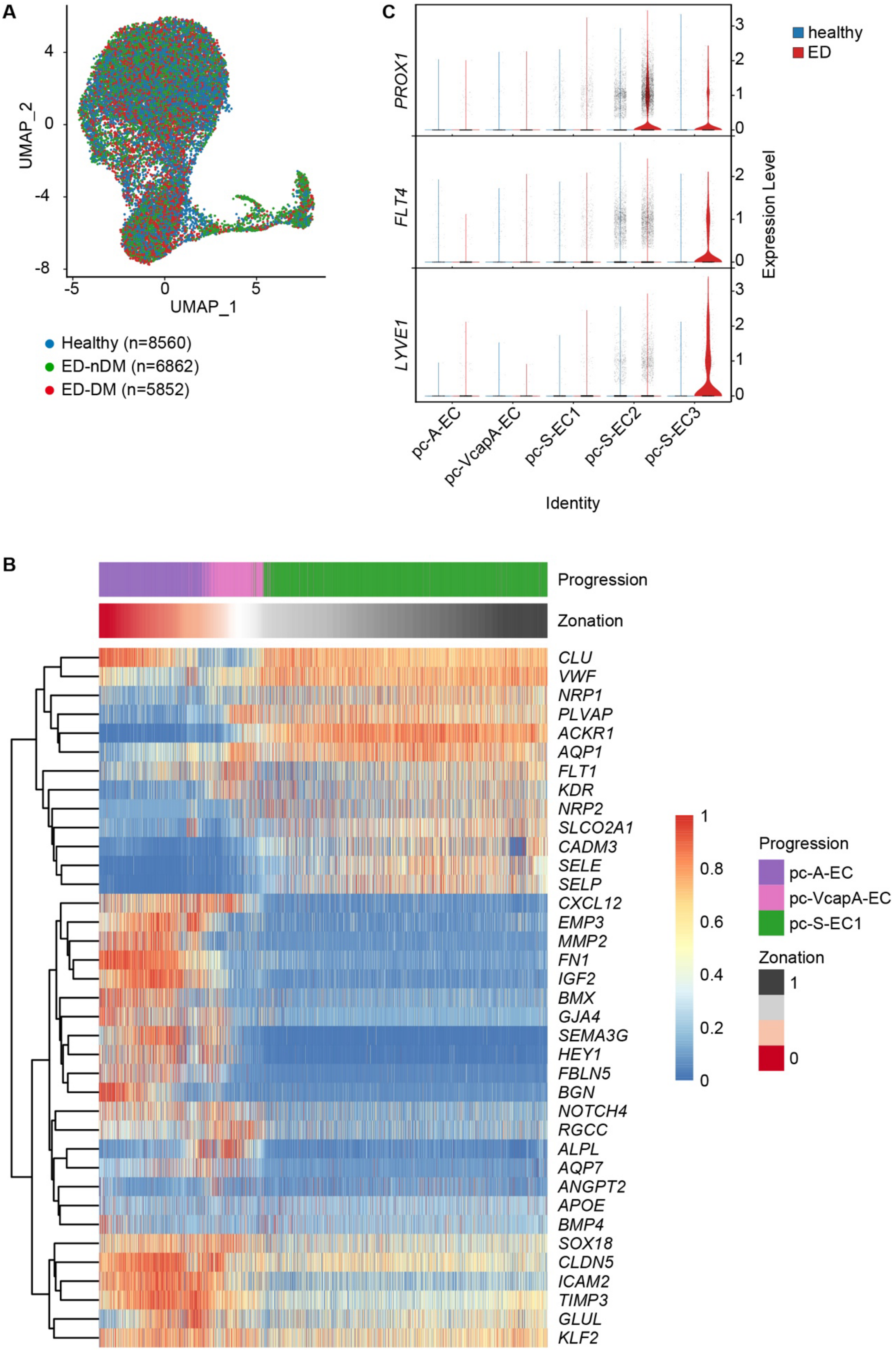
A
UMAP_2
UMAP_1
4
0
-4
-8
-5
0
5
Healthy (n=8560)
ED-nDM (n=6862)
ED-DM (n=5852)
C
PROX1
FLT4
LYVE1
Expression Level
healthy
ED
pc-A-EC
pc-VcapA-EC
pc-S-EC1
pc-S-EC2
pc-S-EC3
Identity
B
Progression
Zonation
CLU
VWF
NRP1
PLVAP
ACKR1
AQP1
FLT1
KDR
NRP2
SLCO2A1
CADM3
SELE
SELP
CXCL12
EMP3
MMP2
FN1
IGF2
BMX
GJA4
SEMA3G
HEY1
FBLN5
BGN
NOTCH4
RGCC
ALPL
AQP7
ANGPT2
APOE
BMP4
SOX18
CLDN5
ICAM2
TIMP3
GLUL
KLF2
1
0.8
0.6
0.4
0.2
0
Progression
pc-A-EC
pc-VcapA-EC
pc-S-EC1
Zonation
1
0

**S2 Fig**

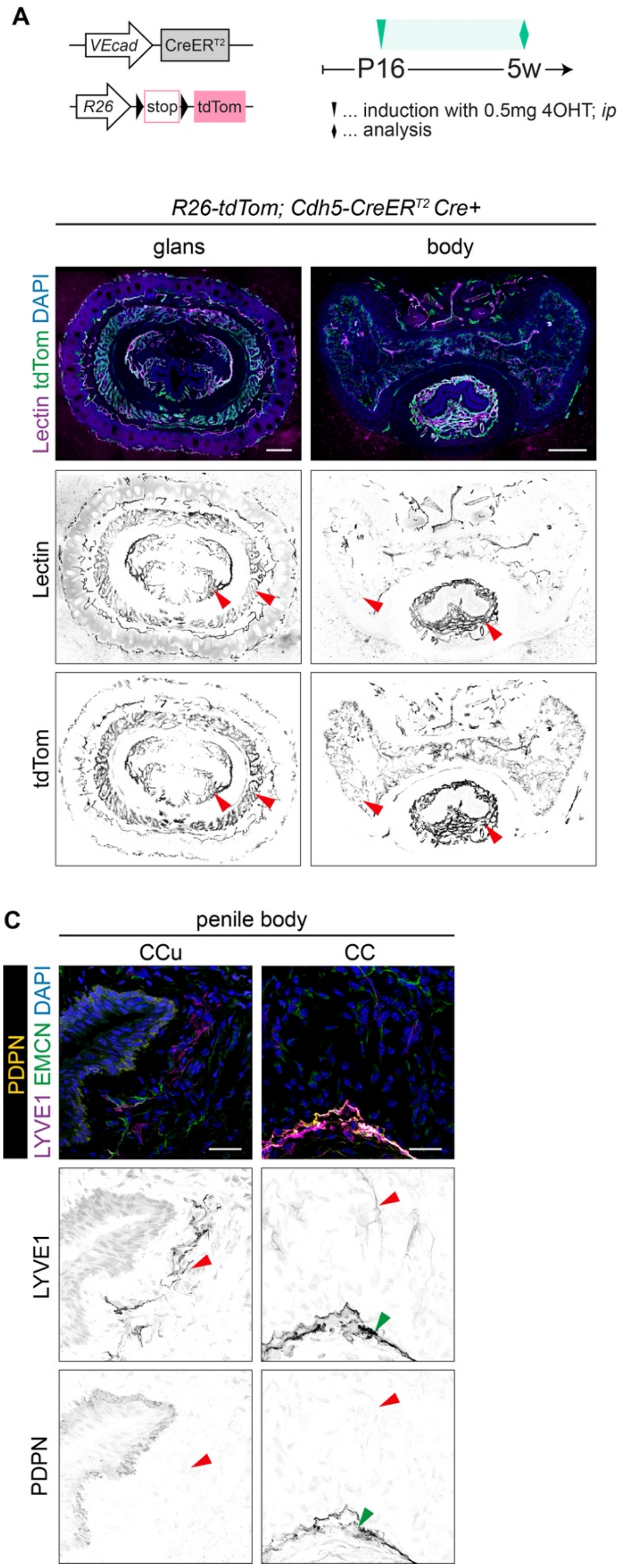
A
VEcad
CreERT2
R26
stop
tdTom
P16
5w
... induction with 0.5mg 4OHT; ip
... analysis
R26-tdTom; Cdh5-CreERT2 Cre+
glans
body
Lectin tdTom DAPI
Lectin
tdTom
B
penile body
CCu
CC
Prox1-GFP Lectin EMCN DAPI
Lectin
Prox1-GFP
EMCN
C
penile body
CCu
CC
PDPN LYVE1 EMCN DAPI
LYVE1
PDPN

### S3 Fig

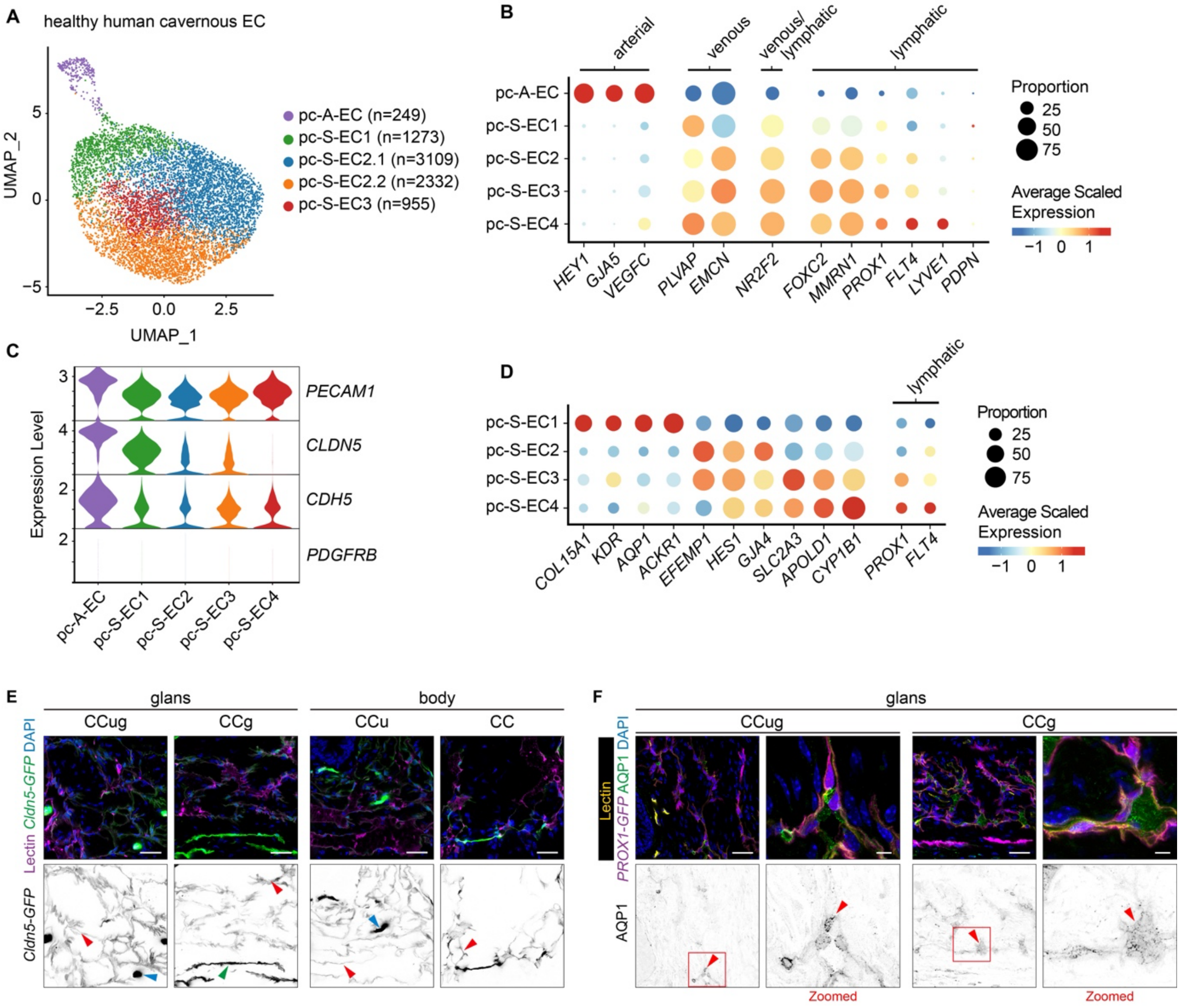
A
healthy human cavernous EC
pc-A-EC (n=249)
pc-S-EC1 (n=1273)
pc-S-EC2.1 (n=3109)
pc-S-EC2.2 (n=2332)
pc-S-EC3 (n=955)
UMAP_1
UMAP_2
B
arterial
venous
venous/ lymphatic
lymphatic
pc-A-EC
pc-S-EC1
pc-S-EC2
pc-S-EC3
pc-S-EC4
HEY1
GJA5
VEGFC
PLVAP
EMCN
NR2F2
FOXC2
MMRN1
PROX1
FLT4
LYVE1
PDPN
Proportion
25
50
75
Average Scaled Expression
C
Expression Level
PECAM1
CLDN5
CDH5
PDGFRB
D
lymphatic
COL15A1
KDR
AQP1
ACKR1
EFEMP1
HES1
GJA4
SLC2A3
APOLD1
CYP1B1
PROX1
FLT4
E
glans
body
CCug
CCg
CCu
CC
Lectin Cldn5-GFP DAPI
Cldn5-GFP
F
glans
CCug
CCg
Lectin
PROX1-GFP AQP1 DAPI
AQP1
Zoomed
Zoomed

**S4 Fig**

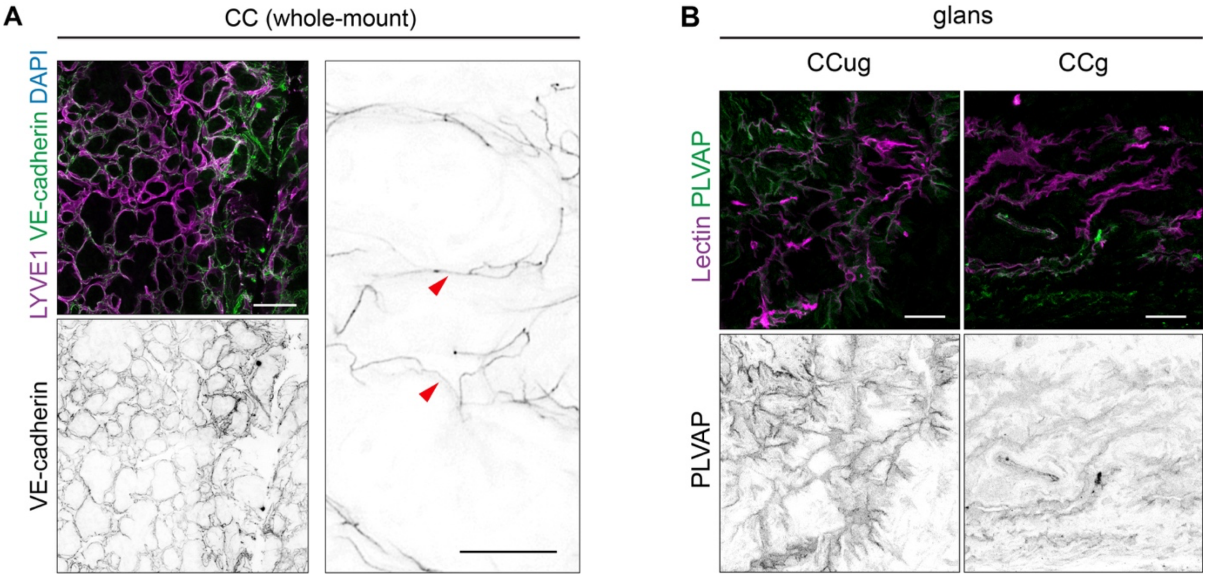

A
CC (whole-mount)
LYVE1 VE-cadherin DAPI
VE-cadherin
B
glans
CCug
CCg
Lectin PLVAP
PLVAP

**S5 Fig**

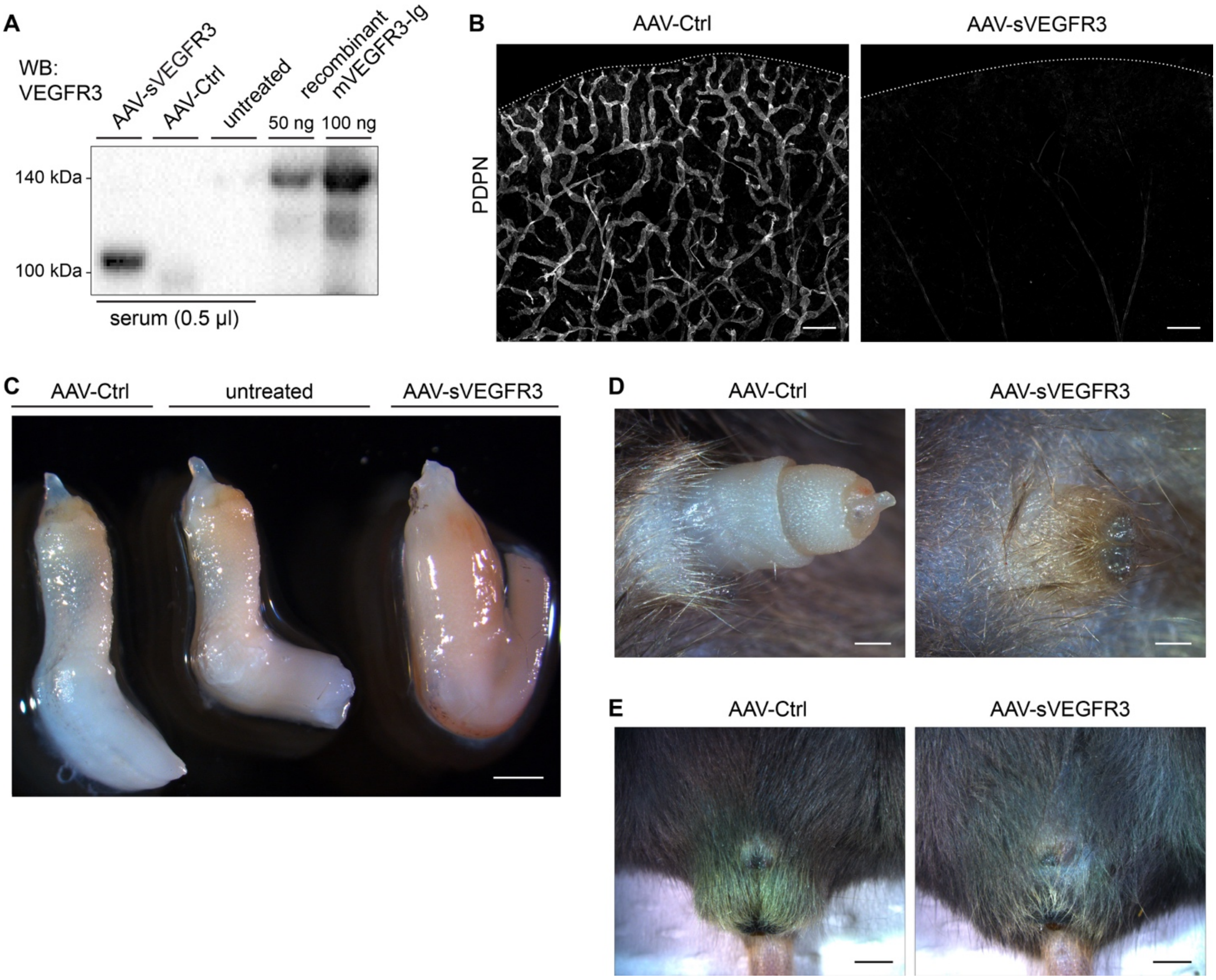
A
WB:
VEGFR3
AAV-sVEGFR3
AAV-Ctrl
untreated
recombinant
mVEGFR3-Ig
50 ng
100 ng
140 kDa
100 kDa
serum (0.5 µl)
B
AAV-Ctrl
AAV-sVEGFR3
PDPN
C
AAV-Ctrl
untreated
AAV-sVEGFR3
D
AAV-Ctrl
AAV-sVEGFR3
E
AAV-Ctrl
AAV-sVEGFR3
